# Controlling protein nanocage assembly with hydrostatic pressure

**DOI:** 10.1101/2020.06.16.154534

**Authors:** Kristian Le Vay, Ben M. Carter, Daniel W. Watkins, T-Y. Dora Tang, Valeska P. Ting, Helmut Cölfen, Robert P. Rambo, Andrew J. Smith, J. L. Ross Anderson, Adam W. Perriman

**Affiliations:** School of Biochemistry, University of Bristol, University Walk, Bristol, BS8 1TD, UK; Bristol Centre for Functional Nanomaterials, HH Wills Physics Laboratory, University of Bristol, Tyndall Avenue, Bristol, BS8 1TL, UK; School of Cellular and Molecular Medicine, University of Bristol, University Walk, Bristol, BS8 1TD, UK; Bristol Composites Institute (ACCIS), University of Bristol, Queen’s Building BS8 1TR, UK; Department of Chemistry, University of Konstanz, Universitätsstraße 10, 78457 Konstanz, Germany; Diamond Light Source Ltd., Diamond House, Harwell Science and Innovation Campus, Fermi Ave, Didcot OX11 0DE, UK; BrisSynBio Synthetic Biology Research Centre, Life Sciences Building, University of Bristol, Tyndall Avenue, Bristol, BS8 1TQ, UK

## Abstract

Controlling the assembly and disassembly of nanoscale protein cages for the capture and internalisation of protein or non-proteinaceous components is fundamentally important to a diverse range of bionanotechnological applications. Here, we study the reversible, pressure-induced dissociation of a natural protein nanocage, *E. coli* bacterioferritin (Bfr), using synchrotron radiation small angle X-ray scattering (SAXS) and circular dichroism (CD). We demonstrate that hydrostatic pressures of 450 MPa are sufficient to completely dissociate the Bfr icositetramer into protein dimers, and the reversibility and kinetics of the reassembly process can be controlled by selecting appropriate buffer conditions. We also demonstrate that the heme B prosthetic group present at the subunit dimer interface influences the stability and pressure lability of the cage, despite its location being discrete from the inter-dimer interface that is key to cage assembly. This indicates a major cage-stabilising role for heme within this family of ferritins.

---

Nanoscale protein cages are attractive scaffolds for bionanotechnology and materials science, where they can be exploited as platforms for constructing robust and configurable therapeutic delivery vectors^1^, vaccines^2^, nanoreactors^3,4^ and templates for the synthesis of diverse nanomaterials^5–8^. These multifunctional containers, both natural^9–13^ and designed^14,15^, offer unparalleled control over size, shape, microenvironment, surface functionalisation and stability when constructing novel bionanomaterials.

The ability to control the assembly of such nanocages is an invaluable tool in the synthesis of complex materials, and can be instrumental in facilitating the encapsulation of non-native nanomaterials. While this can be achieved by exploiting natural^16–18^ or engineered^19–21^ cage metastability, the use of such nanocages could ultimately compromise the robustness of the final assembled material. For nanocages with higher relative stability, harsher environmental conditions^22,23^ are required that can adversely affect the protein cage, its functional modifications or the intended payload for encapsulation. Therefore, new methods are required to circumvent the necessity for harsh chemical conditions or specific interfacial engineering to promote cage instability, and to realise the full potential of these cages in bionanotechnology.

Here, we report on how hydrostatic pressure can be employed to control the disassembly and reassembly of the protein nanocage bacterioferritin from *E. coli* (Bfr). While hydrostatic pressure has been previously employed to dissociate the weakly stable cage-like assembly HSP26^24^, the structure is not truly hollow^25^. There are currently no reports of complete, reversible hydrostatic pressure-induced dissociation in a highly robust nanocage such as ferritin^26,27^. While hydrostatic pressure has been applied to human ferritin to facilitate the loading of doxorubicin and increase protein recovery^27^, the assembly/disassembly of the cage under pressure was not investigated. Specifically, we use synchrotron radiation small angle X-ray scattering (SAXS)^28^ to show that the Bfr nanocage dissociates reversibly under pressure, and that the reassembly can be controlled by altering solution conditions. Hydrostatic pressures of 450 MPa were sufficient to induce reversible dissociation of the Bfr icositetramer into subunit dimers, and the reversibility of the pressure-induced dissociation was found to be highly dependent on ionic strength and temperature, allowing for control of oligomerisation state through pressurisation and selection of buffer conditions. Furthermore, we demonstrate that the pressure lability of the nanocage can be modulated by removal of the native heme B prosthetic group. Our study exploits the ability of SAXS to probe the quaternary, tertiary and secondary structure of proteins, and will provide not only the means for studying the supramolecular assembly of these highly valuable nanocages, but also inform future methodologies for controlling protein nanocage assembly for efficient payload encapsulation.

## Results & Discussion

We initially probed the pressure-induced dissociation of the core-free, or apo-, bacterioferritin icositetramer (ABfr) by gradually raising the applied hydrostatic pressure in a diamond-windowed SAXS pressure cell to 450 MPa in 25 MPa increments, equilibrating for 300 seconds prior to data collection at each pressure. An undulating scattering pattern was observed at low pressures (Figure 1a), characteristic of the hollow nanocage structure. With increasing pressure, fringes corresponding to the hollow sphere form factor broaden and then disappear, indicating increasing polydispersity and decreasing concentration of icositetramer due to dissociation into lower order oligomeric species. The presence of isosbestic points in the reciprocal space and Kratky data (Figure 1a and 1b), indicates that two species contribute to I(*q*) with proportional stoichiometry. At higher pressures (> 300 MPa), a slight drift in the *q* value of the isosbestic points is observed, suggesting additional species likely contribute to the dissociation process.

**Figure 1.**
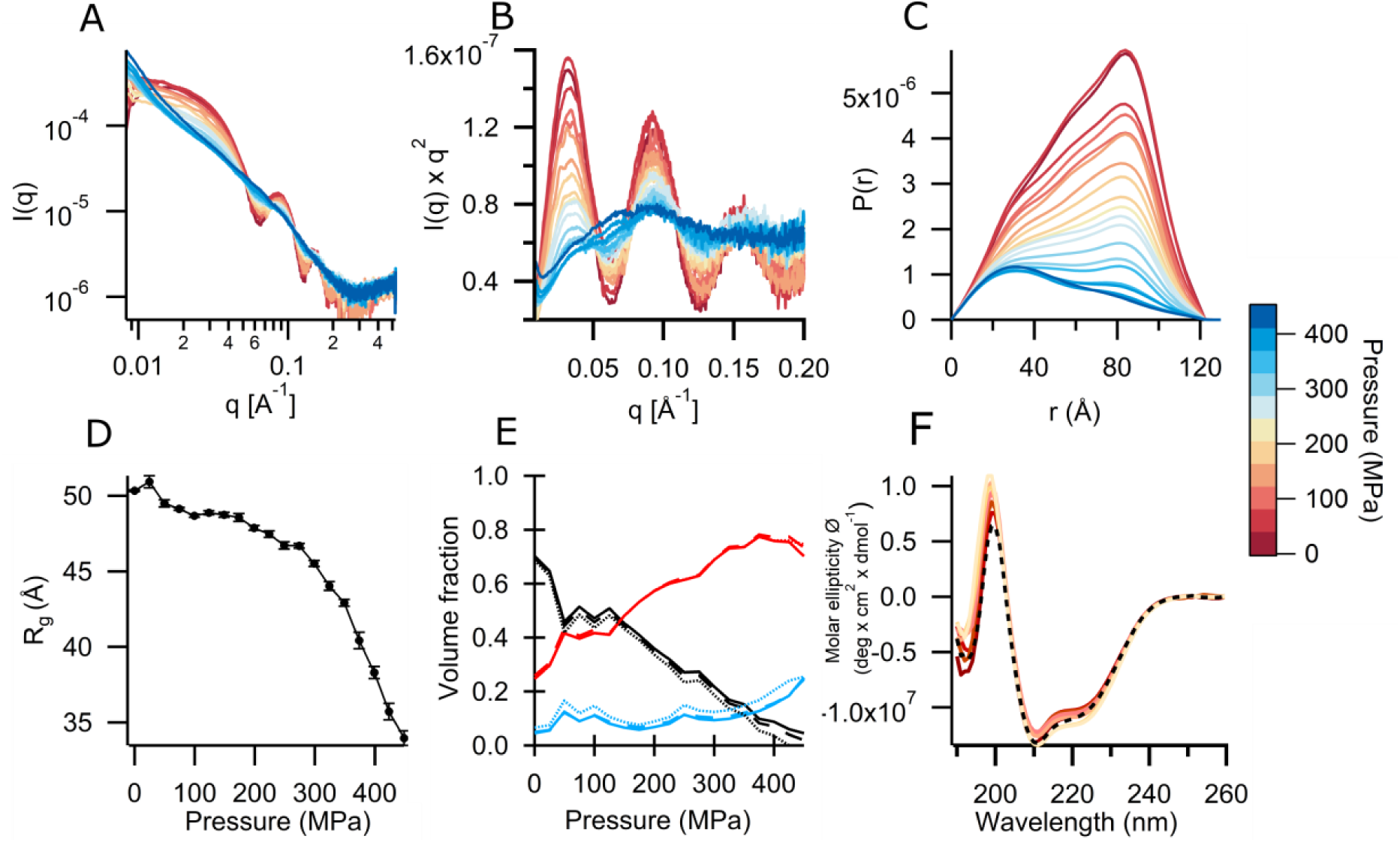
Pressure dissociation of ABfr under equilibrium conditions between 0.1 and 450 MPa measured using high pressure synchrotron radiation SAXS and circular dichroism. **(A)** Reciprocal space SAXS profiles, **(B)** Kratky plots, **(C)** pair distance distribution (P(r)) functions, **(D)** change in radius of gyration, *R*_*g*_, with pressure, **(E)** change in solution oligomer composition with pressure and **(F)** circular dichroism spectra. SAXS data were collected between 0.1 MP (red) and 450 MPa (blue) at a protein concentration of 5 mg mL^-1^ in 45 mM sodium phosphate buffer (pH 7). Real space transformations were performed using BAYESAPP, which uses Bayesian analysis to select parameters such as D_max_ and data noise level.^74^ I(0) and *R*_*g*_ were calculated from real space data. I(0) was normalised to a maximum value of 1. Component volume fraction versus pressure generated from OLIGOMER models of ABfr dissociation. The volume fractions of the icositetramer, *n*-mer and dimer are shown in black, blue and red respectively. The intermediate oligomer states are hexamer (solid line), octamer (dashed line) and dodecamer (dotted line). High-pressure circular dichroism data was collected between 0.1 (red) and 200 MPa (yellow) at a protein concentration of 0.03 mg mL^-1^, 45 mM sodium phosphate, pH 7.

From the pair distance distribution factor (*P(r)*), we determined the radius of gyration, *R*_*g*_ (Figure 1c), which decreases non-linearly from 50 Å to 32 Å over the pressures applied here (Figure 1d). This non-linear change in *R*_*g*_ with pressure is due to the power law dependence of *R*_*g*_ upon number of residues in the scattering particle^29^, rather than suggesting a cooperative dissociation mechanism. The data therefore indicate gradual icositetramer dissociation with multiple intermediates, and a final state in which the majority of species are small subunit oligomers. The *P(r)* distribution of ABfr is characteristic of a hollow sphere until 250 MPa. The maximum of this distribution, *P(r)*_*max*_ (*r* = 84 Å), corresponds to the expected chord length for a hollow sphere and decreases steadily with pressure, indicating gradual loss of the icositetramer. At higher pressures, the magnitude at *r* = 84 Å tends to zero and a new *P(r)*_*max*_ emerges at *r* = 32 Å, corresponding to a subunit oligomer of ABfr previously masked by the fully assembled cage. Importantly, the maximum observed diameter, D_max_, never exceeds the atmospheric pressure value, suggesting that aggregates of intact ABfr particles do not form at higher pressures and that dissociation of the particle into smaller oligomers predominates.

Using a linear combination of theoretical SAXS data from various oligomeric components, we were able to identify possible dissociation pathways^30,31^. Our prediction of possible stable oligomers using the PISA (Proteins, Interfaces, Surfaces and Assemblies)^32^ tool returned only the subunit dimer and icositetramer as stable states (Table S1). We then used single value decomposition (SVD) to determine how many components contribute to the observed variation in the pressure dissociation SAXS data (Figure S1 and S2), and then fitted the dataset using various combinations of oligomer structures derived from the ABfr icositetramer (Table S2, Figure S3 and S4). The SVD analysis indicated two or three major components, with the three most significant accounting for 93.5% of the total contributions (Figure S2). Two component models consisting of the icositetramer and a subunit oligomer provided the worst fits of the dataset (Table S3, Figure S5), though of these, the model consisting of the icositetramer and dimer best represented the data. Three component models generally provided better representations of the dissociation dataset (Figure S5), and all models in which the dimer was the lowest oligomeric state gave the best fit quality. Thus, in this pressure range, we assigned the initial and final state as icositetramer and dimer. Ultimately, we found that the data were best represented by models comprised of icositetramer, dimer and an intermediate hexamer, octamer or dodecamer (Figure 1e). We acknowledge that there is no justification for selecting a specific intermediate state based on these data alone, and therefore refrain from doing so; the equivalence of the models and lack of cooperativity suggests that a range of intermediate states are present during dissociation. Although little work has been carried out on pressure-induced ferritin dissociation, there are many studies detailing pH and denaturant-induced dissociation and reassembly of mammalian ferritin^33–38^. The dissociation products and mechanisms of reassociation observed are consistent with the results obtained here, with dissociation to subunit dimer and reassociation *via* intermediate species including tetramers, hexamers and octamers most commonly reported.

To assess the effect of pressure on the internal structure, secondary structure and folding of ABfr, we used a combination of Kratky analysis and high pressure synchrotron radiation circular dichroism (CD). At low pressure, the Kratky plots for ABfr exhibit multiple peaks that converge to baseline indicative of a globular, spherical particle (Figure 1b). However, at pressures above 375 MPa, the Kratky plots demonstrate a transitioning of ABfr from a spherical particle to a partially folded structure. This may alter tertiary structure and interfacial interactions, destabilising the icositetramer. We observed no significant change in the CD spectra between 0.1 MPa and experimental limit of this technique, 200 MPa (Figure 1f), and predicted structural composition *via* basis spectra remained constant (alpha helix = 0.99, beta sheet, 0.01)^39^. The CD data support the SAXS data in this pressure range, and it is therefore likely that the observed changes in quaternary structure up to 375 MPa are due to the system shifting towards a lower volume state, rather than significant perturbation of the ABfr secondary structure^40^.

We then explored the reversibility of the pressure dissociation process. Initially, ABfr in 45 mM sodium phosphate (NaPi) buffer was pressurised to 450 MPa and held for 5 minutes before pressure release (Figure 2a). An immediate depression in *R*_*g*_ was observed, and the hollow spherical structure was lost after 30 seconds. Fitting the change in icositetramer volume fraction against time with a single exponential function (Figure S6), the observed rate constant (*k*_*diss*_) of dissociation was determined to be 0.114 ± 0.002 s^-1^. Almost complete reassembly occurred over 30 minutes following depressurisation, as apparent from the recovery of the radius of gyration over time (Figure 2a) and the hollow cage form of the P(r) function (Figure 2b). Again, fitting the change in icositetramer volume fraction against time with a single exponential function, the observed rate constant of reassociation (*k*_*ass*_), was determined to be 0.006 ± 0.001 s^-1^ (Figure S6). Following reassociation, we noted that the final *R*_*g*_ and *I(0)* values were slightly lower than initial values (Figure 2a, b). To quantify the degree of reassociation and to control for possible radiation damage or background mismatch, a sample of ABfr in NaPi buffer (45 mM, pH 7) was pressurised under identical conditions without SAXS measurement. Analytical ultracentrifugation (AUC) before pressurisation (Figure 2c, Table S4) demonstrated that ABfr is fully assembled as the icositetramer (S_20, w_ = 15.76, MW = 424 kDa). Following pressurisation (t = 60 minutes), 94.3% of ABfr was assembled in the icositetramer state (S_20, w_ = 16.1, MW = 426 kDa), with the remaining material present as the subunit dimer (S_20, w_ = 3.53, MW = 44.5 kDa). TEM imaging confirmed the presence of assembled ABfr cages both before and after pressurisation, with no apparent change in morphology (Figure 2d, e).

**Figure 2.**
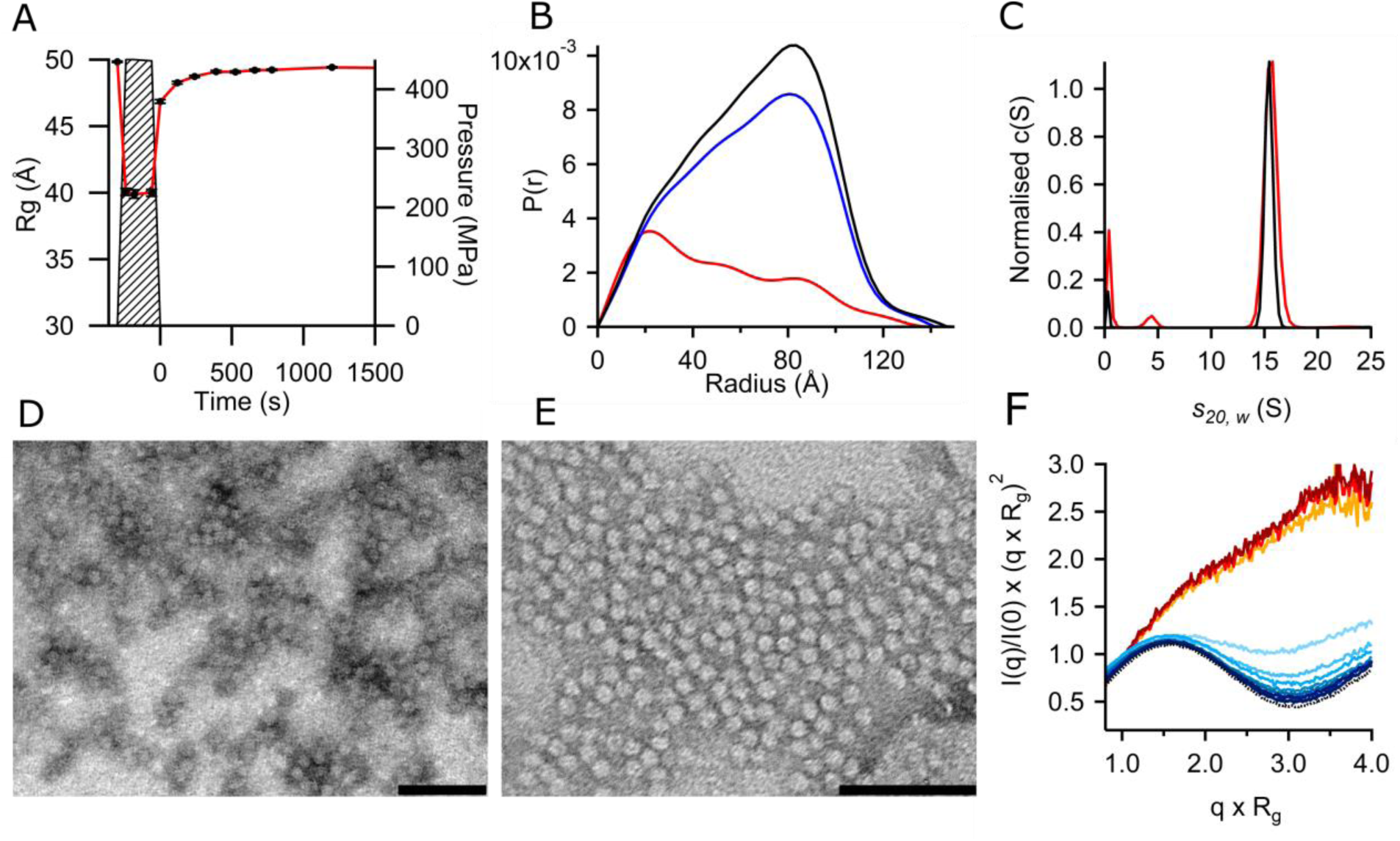
Reassociation of ABfr following pressure induced dissociation measured by SAXS, AUC and TEM. **(A)** Changes in ABfr *R*_*g*_ pre, during and post-pressurization measured by SAXS. Pressure cycle was performed at 25°C NaPi (45 mM, pH 7, I = 123 mM). **(B)** Pair distance distribution (P(r)) functions for ABfr pre- (black), during (red) and post-pressurisation (blue). **(C)** Sedimentation velocity c(S) distribution for ABfr pre-pressurisation (black) and post-pressurisation (red). Sedimentation velocity was measured at a protein concentration of approximately 50 μM in sodium phosphate buffer (45 mM NaPi, 250 mM NaCl, pH 7) using a Beckmann ProteomeLab XL-I at 20°C and 24000 rpm. Data was collected at a wavelength of 418 nm. Data file time stamps were corrected using REDATE, and continuous sedimentation coefficient (c(S)) distributions were fitted using SEDFIT. Buffer density (ρ = 1.003 g cm-3) and viscosity (η = 1.0107 mPa s-1) were measured using an Anton-Paar rolling ball viscometer. **(D)** Negative stain (phosphotungstic acid) TEM images of ABfr pre-pressurisation and **(E)** and post-pressurisation (scale bars = 100 nm). **(F)** Change in normalised Kratky intensity (*I*_*(q=1*.*1)*_/*I*_*(0)*_x(*q*\**R*_*g*_)^2^) with pre, during and post-pressurization. Dotted black trace shows pre-pressurisation data, yellow – red traces show pressurised data at 0, 120 and 300s, light blue – dark blue traces show post pressurisation data with increasing time.

The rate of reassociation under these conditions appears slow, and is significantly slower than the calculated dimer collisional frequency^41^ (*f*_*25*_°_*C*_ = 6.79 x10^5^ S^-1^ at 25°C) demonstrating unequivocally that reassembly is not diffusion limited. To test whether recovery from pressure induced conformational drift influenced the kinetics of reassociation^42,43^, we assessed the degree of protein denaturation by inspection of a normalised Kratky plot (Figure 2f), in which globular protein exhibits a maximum value of 1.104 for qR_g_=√3, whilst unfolded protein has a maximum value of 1.5-2^44^. This plot confirmed that the protein recovers almost immediately from the denatured state after pressure release, so the process is unlikely to be refolding-mediated.

The relatively slow rate of association may also be due to a kinetic barrier in the association process which could arise from repulsive interactions between subunit dimers. To determine the types of interaction dominant in this process, we explored the effect of ionic strength and temperature on the nanocage reassembly, finding strong correlation between these parameters and the rate and completeness of nanocage reassembly (Figure 3, Table S5). We observed the most efficient reassembly in high ionic strength sodium phosphate buffer (45 mM, 250 mM NaCl, pH 7; I = 373 mM)), with higher initial *R*_*g*_ than the low ionic strength buffers and almost complete reassembly at both 5 and 25 °C. In contrast, the assembly was notably slower and less complete in lower ionic strength sodium phosphate buffer (pH 7, 45 mM; I = 123 mM) at both temperatures, with only gradual recovery of *R*_*g*_ over the 1500 seconds of measurement. Most notably, the reassembly process in water was significantly impaired at 25 °C, and was effectively arrested at 5 °C, with no discernable increase in *R*_*g*_ observed over 1500 seconds, indicating the presence of only discrete subunit dimers within this timeframe. Given these data, we propose that the reassembly of ABfr nanocages is likely driven by hydrophobic dispersion forces while opposed by coulombic repulsion.

**Figure 3.**
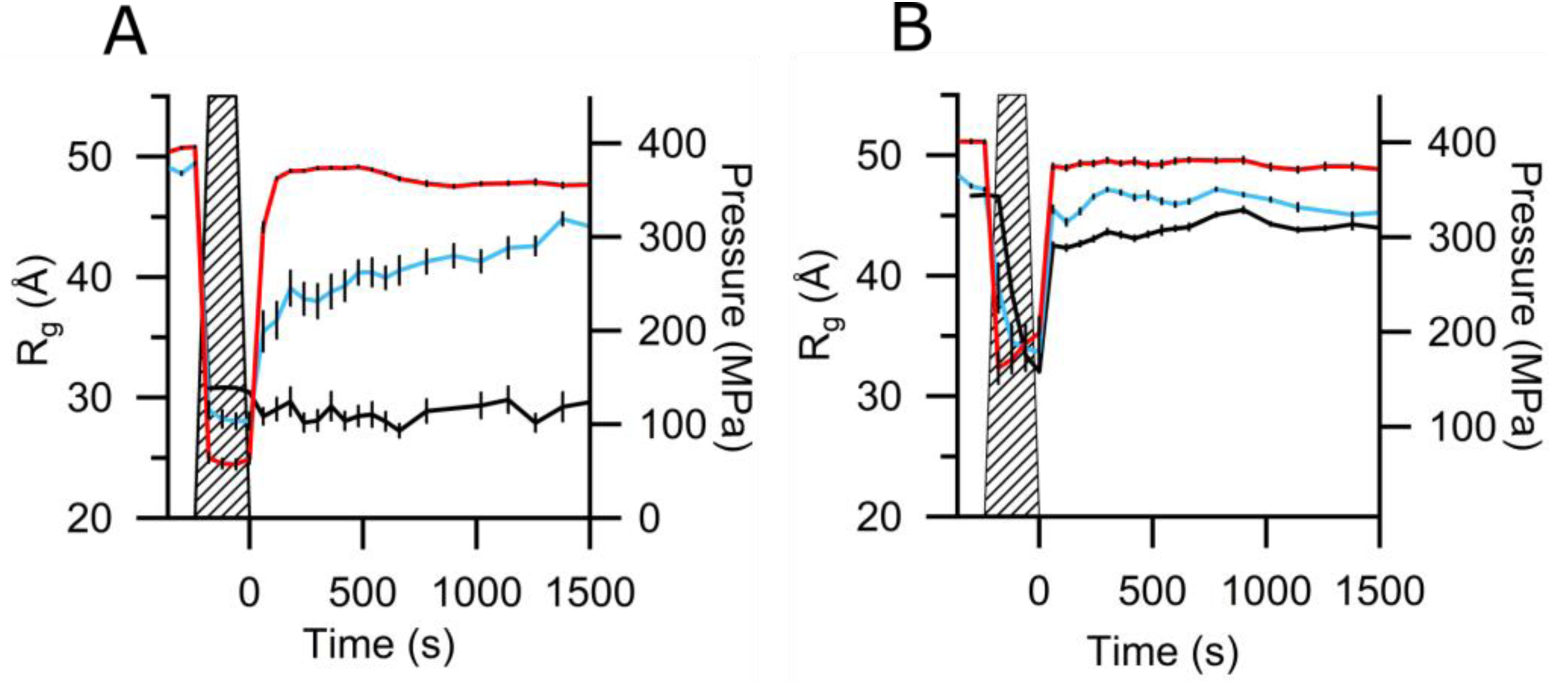
Changes in ABfr *R*_*g*_ pre, during and post-pressurisation. Pressure cycles were performed at 5 °C **(A)** and 25°C **(B)** in H_2_O (I = 0 mM, black), NaPi (45 mM, pH 7, I = 123 mM, blue) and in NaPi (45 mM, pH 7) + NaCl (250 mM) (I = 373 mM, red). Pressure level depicted as hashed region.

Similar ionic strength dependencies on nanocage reassembly were observed for the naturally heme-free *E. coli* ferritin FtnA^45^ following low pH-induced dissociation, and it is likely that the origin of this effect lies at the interface between ferritin subunit dimers, which associate to form the highly charged, carboxylate-rich ion channels at the C3 and C4 interfaces^46–48^. The increased degree and rate of reassembly with temperature is a strong indication that assembly is entropically driven by the formation of weak protein-protein interactions^49^. For oligomeric assemblies, the enthalpic contribution (ΔH) is generally small, because the strength of protein-water and water-water interactions are similar. As such, increasing temperature decreases the Gibb’s free energy of association (ΔG^ass^) and favours assembly.

We also observed a greater degree of dissociation following pressurization at 5°C (5 minutes at 450 MPa) at all ionic strengths tested here; previous studies of pressure-induced viral capsid dissociation have reported similar effects, and were attributed to a strong entropic contribution to the free energy of association^50^. In these cases, higher temperatures lower ΔG^ass^, promoting association in systems where entropic contributions to the free energy dominate. Once the interfacial protein-protein interactions (principally salt bridges and dispersion interactions) are broken and the oligomers dissociate, dipole-dipole protein-water interactions are formed in their place^50,51^. At high pressure, inherently shorter dipole interactions are favoured over dispersion interactions due to the differential effect of compression on bond strength, and dissociation occurs due to the increasing formation of protein-water interactions. In addition, the hydration of hydrophobic surfaces is more favourable at high pressure, as the ordered solvation shell is denser than the bulk solvent. Both similar and contrasting behaviours have been reported for viral capsids under pressure. Silva *et al*.^52,53^ demonstrated that the 86-subunit bromegrass mosaic virus capsid undergoes a reversible partial dissociation into dimers upon application of pressure (10*%* dissociation at 200 MPa). In contrast, the turnip yellow mosaic virus irreversibly decapsidates rather than dissociates under pressure, resulting loss of RNA and formation of a holed capsid, as the subunit interface contains few pressure sensitive salt bridge and is rich in pressure insensitive hydrogen bonding interactions^54^.

We subsequently investigated the ability of the native heme B prosthetic group to modulate cage stability. Using a well-established procedure for tetrapyrrole extraction, we removed heme B using an acidified water:2-butanone mixture and confirmed the successful removal by UV/visible spectroscopy (Figure 4a)^55^. We next explored the composition of oligomeric species in this apo-apo-Bfr (AABfr) by sedimentation velocity analytical ultracentrifugation (SV-AUC, Figure 4b). In contrast to the heme containing ABfr, the SV-AUC distribution reveals a mixture of assembled icositetramer (S_(20, w)_ = 15.85, MW = 427.1 kDa, 62.1%) and subunit dimer (S_(20, w)_ = 2.97, MW = 34.7 kDa, 37.9%). The icositetramer peak is broadened, indicating greater polydispersity and potentially incomplete assembly. We further characterized this mixture of species by high-performance liquid chromatography-SAXS (HPLC-SAXS) (Figure 4c), and observed scattering patterns consistent with both the assembled icositetrameric nanocage (*R*_*g*_ = 49.88 Å), and a smaller ellipsoidal or parallelepipedal particle^56^. The extrapolated *R*_*g*_ (22.21 Å) of the smaller particle is in good agreement with that calculated for the subunit dimer (PDBID: 2VXI, *R*_*g*_ = 21.16 Å)^57^, and the Kratky plot (Figure 4d) indicates a compact structure, although with a greater degree of disorder than the AABfr icositetramer. We used bead modelling to analyse the SAXS data for both the ABfr and AABfr icositetramers and found excellent agreement between the hollow spherical models and the published Bfr crystal structure (Figure 4e)^57^. Similarly, the bead model of the AABfr subunit dimer closely overlays with the crystal structure of the AABfr dimer (PDBID: 4CVP)^58^. To further probe the effects of removing heme B from the nanocages, we used circular dichroism spectroscopy to determine the thermal stabilities of the Bfr samples. We found the denaturation midpoints of ABfr (T_m_ = 68°C) and AABfr (T_m_ = 58°C) in agreement with previously reported values^59,60^, confirming that removal of heme B has an overall destabilizing effect on the protein. Comparison of the ABfr and AAbfr subunit dimer crystal structures reveals slight secondary and tertiary structure differences that may impact the ability of the dimer to assemble into the icositetramer (Figure S7). In particular, the position of the E helix is shifted in the heme free protein; this helix lies at the tetramerization interface and is essential for cage assembly (Figure S8)^61^.

**Figure 4.**
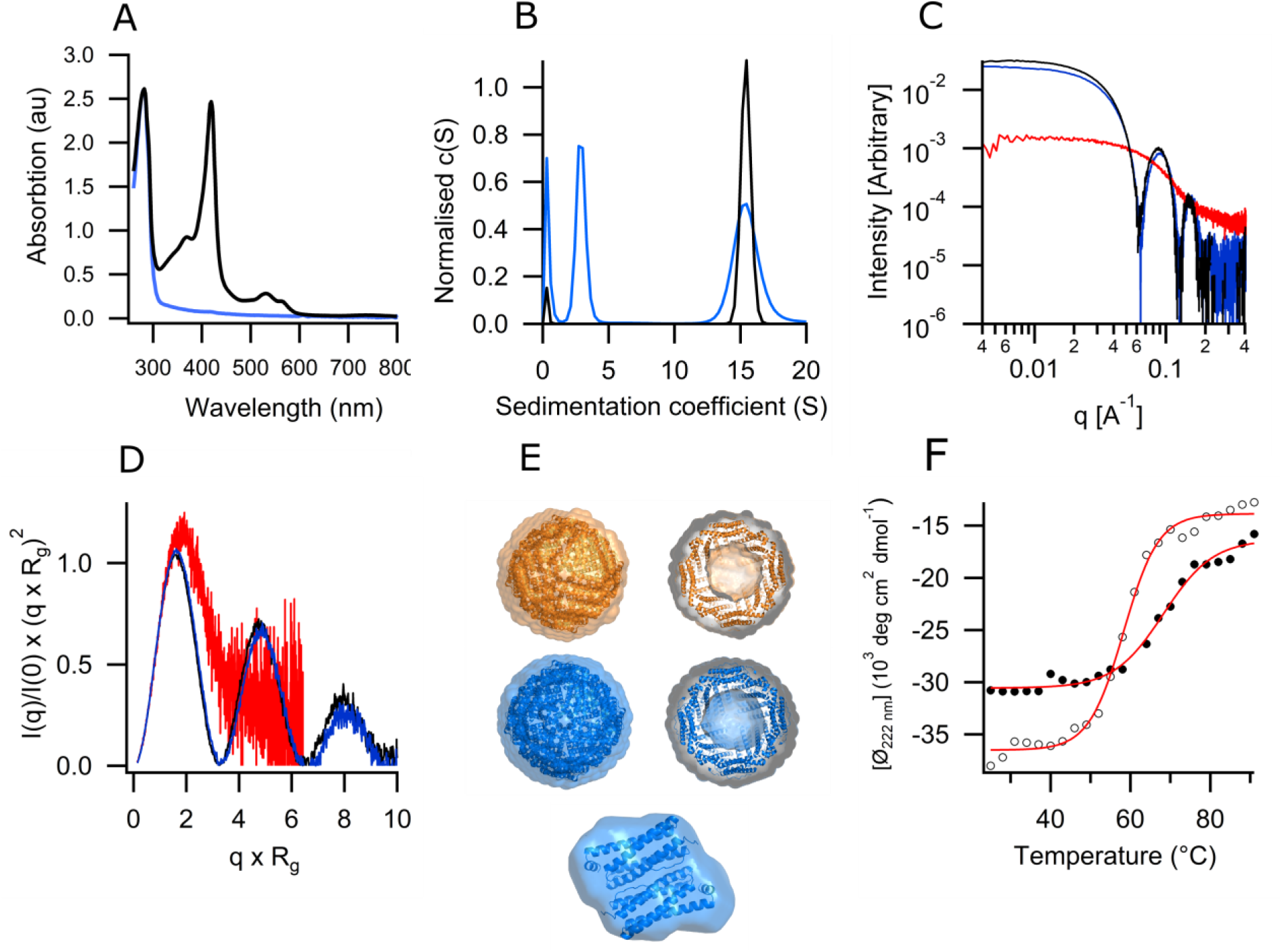
Physicochemical characterization of ABfr and AABfr under ambient pressure. **(A)** UV/visible spectra of ABfr (black) and AABfr (blue). UV-visible spectra were recorded measured at a protein concentration of approximately 50 μM in sodium phosphate buffer (45 mM NaPi, 250 mM NaCl, pH 7). Curves are normalised to A280 = 1 to highlight differences in heme absorbance. **(B)** SV-AUC c(s) distributions for ABfr (black) and AABR (blue). Sedimentation velocity was measured at a protein concentration of approximately 50 μM in potassium phosphate buffer (45 mM KPi, 250 mM NaCl, pH 7) using a Beckmann ProteomeLab XL-I at 20°C and 24000 rpm. Data was collected at a wavelength of 280 nm. Data file time stamps were corrected using REDATE, and continuous sedimentation coefficient (c(S)) distributions were fitted using SEDFIT. Buffer density (*ρ* = 1.003 g cm^-3^) and viscosity (*η* = 1.0107 mPa 865-) were measured using an Anton-Paar rolling ball viscometer. The protein partial specific volume was calculated from the primary sequence (*ṽ*= 0.736 cm^3^ g^-1^). **(C)** HPLC-SAXS profiles and **(D)** Kratky plots for ABfr icositetramer (black) and AABfr icositetramer (blue) and dimer (red). **(E)** *Ab initio* bead model of ABfr (icositetramer, orange) and AABfr (icositetramer and dimer, blue), overlaid with corresponding crystal structures (2VXI and 4CVP)^57,58^. Real space transformations were performed using ScÅtter. The maximum diameter was determined by selecting values that resulted in high reciprocal fit quality, and produced smooth, oscillation-free, real-space functions that decreased smoothly to zero at high radius. A constant background was applied in the transformation, and real space distributions were refined using the L1 norm of the Moore coefficients as a regularisation target.The *ab initio* models were produced from refined pair distance distribution functions using DAMMIF, and are DAMAVER averages of 23 runs. The models and crystal structures were visualized using PyMOL. **(F)** Thermal denaturation far-UV circular dichroism spectra and fits for ABfr (open circles) and AABfr (black circles). Data were collected at B23, Diamond Light Source, UK. Measurements were performed in potassium phosphate buffer (45 mM, pH 7) at a protein concetnration of 0.5 g L^-1^. Raw data was converted to mean residue elipticity and secondary structure analysis was performed using CAPITO.

These data indicate that the binding of heme B not only leads to an increase in thermal stability of the bulk Bfr protein, but it specifically stabilises the nanocage assembly. While such increases in thermal stability induced through cofactor binding in heme-proteins are widely reported, the bound cofactor’s impact on cage stability and assembly is unknown^62^. To determine the effect of heme B on the pressure stability of Bfr, we pressurised AABfr to 450 MPa in 25 MPa increments, allowing time at each step for equilibration before SAXS measurement (Figure 5a). We found the real space distribution for ABfr at ambient pressure is representative of a mixture of assembled icositetramers and smaller subunit oligomers, in good agreement with the SV-AUC data described above (Figure 5b). The icositetramer nanocage structure visible in the real space distribution is rapidly lost with applied pressure, whilst the final state resembles a mixture of the subunit dimer and other lower order oligomers. The Kratky plots demonstrate a less globular structure than ABfr at similar pressures, and do not reveal any further unfolding over the pressure range (Figure 5c). The initial *R*_*g*_ for AABfr (42 Å) Is notably lower than that of ABfr (50 Å) due to the mixture of icositetramer and dimer at ambient pressure. The *R*_*g*_ remains constant until 175 MPa, then deceases, reaching a plateau at 350 MPa, demonstrating that AABfr is significantly less pressure stable than ABfr (Figure 5d). The final *R*_*g*_ of 35 Å, was higher than that of ABfr (32 Å), suggesting a dissociation endpoint with a different oligomeric state.

**Figure 5.**
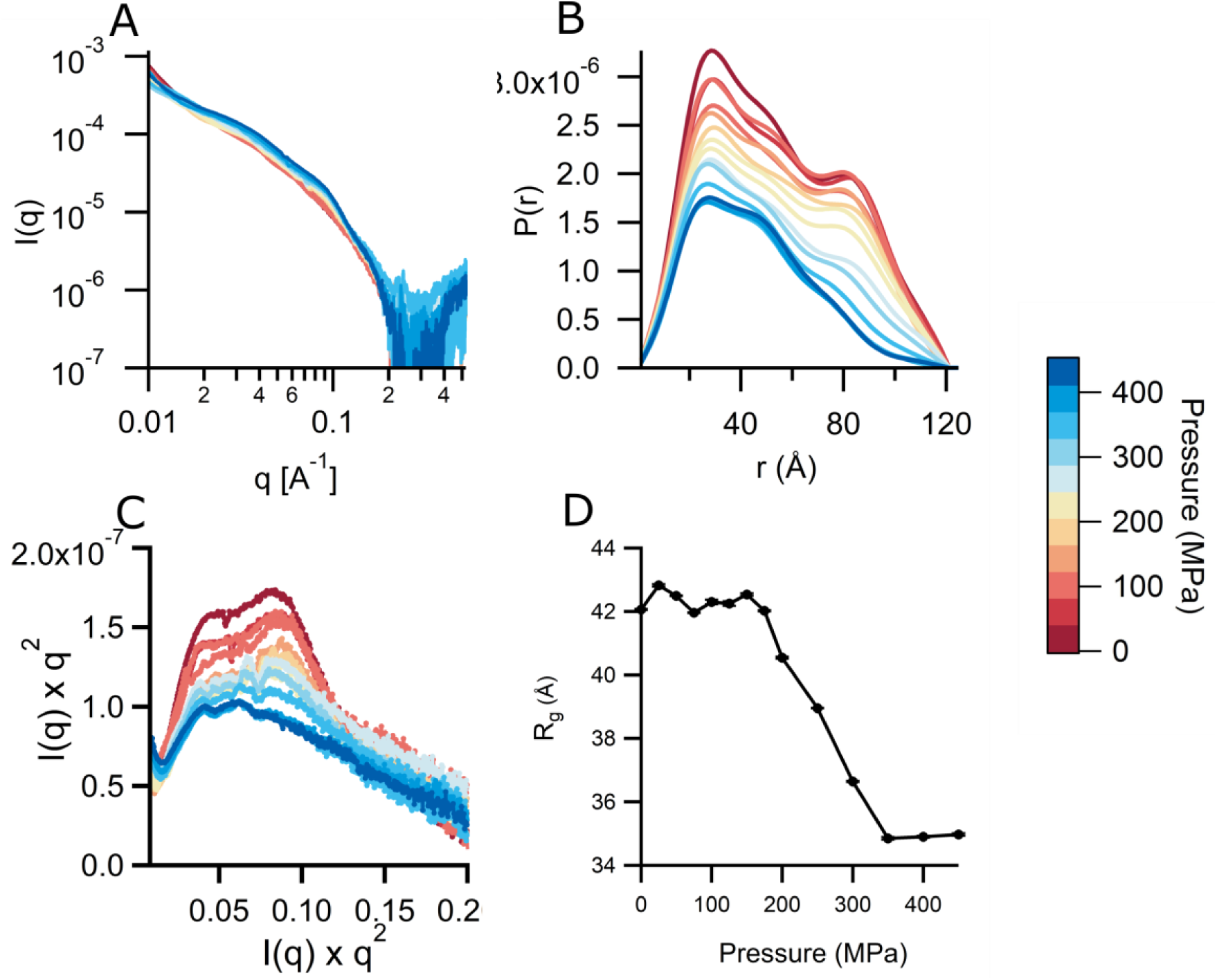
Pressure dissociation of AABfr under equilibrium conditions between 0.1 and 450 MPa. **(A)** reciprocal space SAXS profiles, **(B)** Pair distance distribution (P(r)) functions, **(C)** Kratky plots and **(D)** Radius of gyration, *R*_*g*_. SAXS data was collected between 0.1 MP (red) and 450 MPa (blue) at a protein concentration of 5 mg mL^-1^ in 45 mM sodium phosphate buffer (pH 7). Real space transformations were performed using BAYESAPP, which uses Bayesian analysis to select parameters such as D_max_ and data noise level.^74^ *R*_*g*_ was calculated from real space data.

## Conclusions

We have demonstrated here that hydrostatic pressure is a valuable method to control and modulate the assembly, disassembly and oligomeric composition of the bacterioferritin nanocage. It is highly likely that this methodology can be extended to other protein nanocages, especially those stabilized primarily by hydrophobic interactions. Since the method we report here is also gentle, tunable and not limited to intrinsically metastable or mutationally compromised cages, more robust hybrid materials tolerant of harsh environmental conditions are potentially accessible. Full cage dissociation might also provide a route to higher loading ratios of therapeutic molecules or larger payloads unable to traverse the protein cage, leading, for example, to significantly improved synthetic routes to nanocage-based drug delivery vehicles.

Furthermore, we have also demonstrated that the heme B prosthetic groups significantly enhance the Bfr nanocage stability. While it has been previously reported that heme B facilitates electron transfer and iron release from the Bfr core, this can occur in the absence of the prosthetic group and it is notable that heme B is absent in many/most of the known ferritins. It therefore seems plausible that an additional major role of heme B in Bfr is to stabilise the protein nanocage, thus enabling the retention of the protein superstructure for iron mineralisation.

## Methods

### Protein expression and purification

Bacterioferritin expression was carried out in T7 express *E. coli* BL21 (DE3) cells using a modified pUC119 plasmid, PGS281 as described in Andrews *et al*^63^. Cultures were grown aerobically at 37 °C in LB media containing 34 µg mL^-1^ carbenicillin. Flasks were shaken for 24 hours at 200 rpm, before cultures were harvested by centrifugation (10 minutes, 7277 ×g rcf). The pellets were washed then re-suspended in lysis buffer (1.5 mM KH_2_PO_4_, 8 mM Na_2_HPO_4_, 150 mM NaCl, 3 mM KCl, pH 7). Phenylmethylsulfonyl fluoride (PMSF) (1 mM) was added, and then cells were lysed using a probe sonicator (3 × 20s, maximum amplitude). The crude extract was then centrifuged (47808 ×g, 60 minutes). The supernatant was decanted, heated to 70 °C for 15 minutes, then cooled and centrifuged (47808 ×g, 30 minutes). The supernatant was concentrated using a centrifugal concentrator (MWCO = 50 kDa) to half the original volume. The extract was purified by size exclusion chromatography using a Sephadex S200 26/600 column equilibrated with lysis buffer at a flow rate of 2.3 mL *per* minute. Bfr containing fractions were pooled, concentrated, and then further purified by anion exchange chromatography using a Q-Sepharose FF column. The target protein was eluted with a linear gradient of 0-0.3 M NaCl in histidine buffer (20 mM histidine.HCl, pH 5.5). The purified protein was dialysed in a solution of EDTA (10 mM) and DTT (5 mM) to remove the native core, forming ABfr.

### High pressure SAXS measurement

Data collection was carried out at the I22 beamline, Diamond Light Source (DLS, Harwell, UK), using a hydrostatic pressure cell with a maximum operation pressure of 500 MPa^28^. Samples were loaded into thin-wall polycarbonate capillaries (θ=2 mm), which were then sealed with a rubber bung and two-part adhesive. The beam energy was 18 keV (λ = 0.69 Å) to reduce absorption from the diamond windows and water in the beam path. ABfr samples were prepared at a concentration of 5 mg mL^-1^ in a range of buffers and centrifuged (10 minutes, 16000 ×g) prior to measurement. The collection time for individual measurements was between 6 and 60 s. Hydrostatic pressure experiments were performed under both equilibrium and dynamic, time resolved conditions. In the former, the pressure was raised in incremental steps allowing time for equilibrium before measurement. The beam was blocked during equilibration to prevent radiation damage. In dynamic experiments the sample was pressurised, held at pressure, then returned to ambient pressure whilst measuring the time resolved scattering pattern. Pressure jump experiments were conducted in which pressure was increased rapidly to 450 MPa whilst measuring the time resolved scattering. The data were collected using a Pilatus P3-2M detector. The sample to detector distance was 6 m, providing a *q*-range of 0.008-0.52 Å^-1^. The two-dimensional data sets were reduced using DAWN^64^. Briefly, the *q-*axis was calibrated using with a silver behenate standard data, then detector images were masked, radially averaged from the beam centre and normalised to absolute intensity using a glassy carbon standard. Background subtraction was carried out using a user-written python script (Figure S9 and S10) to account for background mismatch in the high-*q* data region. This mismatch resulted from the necessity of using different capillaries for sample and background, and slight changes in cell and capillary position when changing samples. Whilst this mismatch was generally low, systemic under-subtraction at low pressure and over-subtraction at high pressure was present across data sets. The background measurement was multiplied by a constant to align the high-*q* region with that of the sample. The median I(*q*) of the region *q* = 0.3-0.4 Å^-1^ was determined for both sample and background data. The background was then multiplied by a constant, *k*, such that the corrected median I(*q*) of the background was equal to the median I(*q*) of the sample minus a small constant, *A*. The scaling constant was therefore:

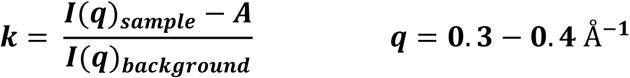

This method was effective in background matching the high-*q* data region and preventing over and under subtraction with *k* = 1 × 10^−5^. Example data corrections for ABfr at 0.1 MPa and 200 MPa are shown in Figure S9. The background adjustment results in lower overall I(*q*) relative to the unprocessed data, but the shape of the scattering curve is preserved. Artefacts resulting from Kossel lines are visible in the data at 200 MPa at *q* = 0.016 and 0.7 Å^-1^ despite background subtraction. The effect is most visible in the 2D detector image, shown in Figure S10. Kossel lines are observed between 100-300 MPa in both background and sample images. The intensity of the effect is pressure dependent, and line position is dependent on the orientation of the diamond windows. Care was taken to repeat background measurements following window changes. However, mismatched line intensity between sample and background resulted in artefacts in the reduced data. The mismatch occurs due to the pressure dependence of line intensity: whilst pressure control is accurate to 0.1 MPa in this apparatus, slight pressure loss often occurs during measurement due to leakage. Consequently, pressure differences between samples and background measurements of up to 0.3 MPa are observed. Although visible in the data, the artefacts are small and not expected to affect subsequent fitting and data analysis. Parasitic scattering is present in some data sets, particularly where Kossel lines intersect with the beam-stop (Figure S10 (200 MPa)). This effect is not effectively removed by background subtraction, and manifests as aggregation-like scattering at *q* < 0.04 Å^-1^, which precludes the use of Guinier methods for the approximation of I(0) and *R*_*g*_, but does not affect the determination of these parameters from the indirect Fourier transform of the data.

### High pressure liquid chromatography small angle X-ray scattering

HPLC-SAXS measurements were carried out using the B21 beamline at the Diamond Light Source, Harwell, UK. Measurements were taken with a camera length of 3.9 m and a beam energy of 12.4 keV. The beam cross section was 1 × 5 mm. The scattering angle and absolute intensity were pre-calibrated using silver behenate and glassy carbon standards. Samples were measured in a glass capillary at a concentration of 2-5 mg mL^-1^ and a temperature of 25°C. Data were collected in 30 successive frames of one second each. Inline HPLC separation was carried out using a Shodex KW 403 column. The 2D SAXS patterns obtained were radially averaged from the beam centre, normalised to an absolute intensity scale using a glassy carbon standard and corrected for background scattering from buffer components. ScÅtter was used to determine basic SAXS invariants and determine the pair distance distribution (P(r)) functions from the experimental data as described previously^65,66^. Ab initio modelling was carried out using DAMMIF and DAMAVER with no symmetry restrictions^67^. The crystallographic model of *E. coli* bacterioferritin was obtained from the Protein Data Bank (PDB code: 2VXI, 4CVP), and aligned to the reconstructed structural models using the program SUPALM^68^.

### Circular Dichroism

Circular dichroism (CD) spectroscopy was used to determine the relative thermal stabilities of ABfr and AABfr and to investigate the secondary structure of both proteins. Solutions of ABfr or AABfr (0.5 g L^-1^) were prepared in 45 mM sodium phosphate buffer (pH 7). The far UV spectrum (190-260 nm) was measured at 25 °C, then the temperature was increased in 3 degree increments allowing for equilibration (300s) at each step. The helical content was measured by monitoring mean residue ellipticity at λ= 222 nm. The thermal denaturation midpoint temperature was determined using sigmoidal fits of the raw the [θ_222 nm_] data and from the maxima in first differential of the [θ_222 nm_] data (Figure 4f). Secondary structure composition was determined by linear combination of basis spectra using CAPITO^39^.

### High pressure circular dichroism

High pressure circular dichroism measurements were carried out at the B23 beamline, Diamond Light Source (DLS, Harwell, UK), using a hydrostatic pressure cell with a maximum operation pressure of 200 MPa^69^. Results obtained were processed using CDApps and OriginLab. Secondary structure estimation from CD spectra was carried out using the CAPITO CD analysis and plotting tool^39^.

### Sedimentation velocity analytical ultracentrifugation

Sedimentation velocity was measured in sodium phosphate buffer (45 mM NaPi, 250 mM NaCl, pH 7) using a Beckmann ProteomeLab XL-I at 20 ° C and 24000 rpm. Data were collected at a wavelength of 418 nm. Date file time stamps were corrected using REDATE, and continuous sedimentation velocity coefficient (c(S)) distributions were produced using SEDFIT. Buffer density and viscosity were measured using an Anton-Paar rolling ball viscometer. The protein partial specific volume was calculated from the primary sequence (*ṽ*= 0.736 cm^3^ g^-1^).

### Transmission electron microscopy

All transmission electron microscopy data were collected by using a JEOL 1200 MK2 TEM at the University of Bristol. Bfr samples were deposited onto carbon film coated copper mesh grids and stained using phosphotungstic acid, which is not internalised by the protein cage.

### PISA analysis of ABfr

A bioinformatics approach was adopted in order to identify potential dissociation states of ABfr under pressure. The Proteins, Interfaces, Structures and Assemblies (PISA) tool was used to calculate overall solvent accessible surface areas, solvation free energies (ΔG^int^) and dissociation free energies (ΔG^diss^) for stable subunit oligomers of ABfr^32,70,71^. The PISA analysis identifies thermodynamically stable assemblies of subunits based on interfacial interactions. For ABfr, only the fully assembled icositetramer and dimer were identified as stable quaternary structures in solution. The surface areas and free energies of these species are shown in Table S1. The free energy of dissociation, ΔG^diss^, corresponds to the free energy difference between associated and dissociated states. A positive value indicates that the assembled state is stable under standard conditions.

### SVD analysis and oligomer model fitting of ABfr pressure dissociation

SVDPLOT and Ultrascan II were used to produce a set of basis eigenvectors and eigenvalues for the ABfr dataset (Figure S1)^31,72^. A non-parameterised runs test was used to identify non-random curves in the eigenvector set^73^. Eigenvectors with p < 0.05 were deemed to be non-random, yielding 12 significant eigenvectors. Inspection of the ABfr eigenvectors revealed that those with the highest three eigenvalues are smooth curves reminiscent of scattering form factors, and so are likely to contain structural information corresponding to the initial icositetramer and oligomeric dissociation products. The subsequent curves are structured, but contain significant noise, and may contain contributions from minor species as well as background components due to path length variation an imperfect background subtraction. The data were then reconstructed incrementally by adding eigenvectors to the model in order of decreasing significance. The root mean squared deviation between the reconstructed datasets and the experimental curves was calculated at each stage (Figure S2).

Subunit oligomer models were extracted from the crystal structure of the full ABfr icositetramer (2VXI), and theoretical scattering data from these models was calculated using CRYSOL (Figures S3 and S4)^30^. OLIGOMER was then used to fit linear combinations of these oligomers to the experimental dataset^31^. Fit quality was assessed by the reduced chi-squared value and by inspection of fit residuals (Figure S5 and Table S3).

## Acknowledgements

This work was supported at the University of Bristol by the Bristol Centre for Functional Nanomaterials (EPSRC Doctoral Training Centre Grant EP/G036780/1) through a studentship for K.L.V. We acknowledge Diamond Light Source for time on I22 under proposals SM8237, SM9367, and SM11615, for time on B21 under proposals SM10054, SM1318 and SM16020, and for time on B23 under proposal SM14069.

## Author Contributions

J.L.R.A and A.W.P conceived the project; K.L.V., D.W., B.C., D.T., V.T., and A.J.S. performed the experiments; K.L.V., B.C., H.C., R.R., A.J.S., J.L.R.A. and A.W.P. discussed the results; K.L.V. and J.L.R.A. wrote the manuscript.

## Supporting Information

PISA analysis, detailed AUC data, structural models and additional SAXS data including single value decomposition, calculated P(r) distributions, OLIGOMER models, ABfr dissociation data and background subtraction examples are provided in the supporting information.

## References

1. Bhaskar, S. & Lim, S. Engineering protein nanocages as carriers for biomedical applications. NPG Asia Materials 9, e371–e371 (2017).

2. Kanekiyo, M. et al. Self-assembling influenza nanoparticle vaccines elicit broadly neutralizing H1N1 antibodies. Nature 499, 102–106 (2013).

3. Ren, H., Zhu, S. & Zheng, G. Nanoreactor design based on self-assembling protein nanocages. Int. J. Mol. Sci. 20, (2019).

4. Maity, B., Fujita, K. & Ueno, T. Use of the confined spaces of apo-ferritin and virus capsids as nanoreactors for catalytic reactions. Curr. Opin. Chem. Biol. 25, 88–97 (2015).

5. Douglas, T. & Stark, V. T. Nanophase cobalt oxyhydroxide mineral synthesized within the protein cage of ferritin. Inorg. Chem. 39, 1828–1830 (2000).

6. Yamashita, I., Hayashi, J. & Hara, M. Bio-template Synthesis of Uniform CdSe Nanoparticles Using Cage-shaped Protein, Apoferritin. Chem. Lett. 33, 1158–1159 (2004).

7. Ueno, T. et al. Size-selective olefin hydrogenation by a Pd nanocluster provided in an apo-ferritin cage. Angew. Chemie - Int. Ed. 43, 2527–2530 (2004).

8. Douglas, T. et al. Protein engineering of a viral cage for constrained nanomaterials synthesis. Adv. Mater. 14, 415–418 (2002).

9. He, D. & Marles-Wright, J. Ferritin family proteins and their use in bionanotechnology. N. Biotechnol. 32, 651–657 (2015).

10. Jutz, G., Van Rijn, P., Santos Miranda, B. & Böker, A. Ferritin: A versatile building block for bionanotechnology. Chem. Rev. 115, 1653–1701 (2015).

11. Putri, R. M. et al. Structural Characterization of Native and Modified Encapsulins as Nanoplatforms for in Vitro Catalysis and Cellular Uptake. ACS Nano 11, 12796–12804 (2017).

12. Choi, S. H., Kwon, I. C., Hwang, K. Y., Kim, I. S. & Ahn, H. J. Small heat shock protein as a multifunctional scaffold: Integrated tumor targeting and caspase imaging within a single cage. Biomacromolecules 12, 3099–3106 (2011).

13. Azuma, Y., Edwardson, T. G. W. & Hilvert, D. Tailoring lumazine synthase assemblies for bionanotechnology. Chemical Society Reviews 47, 3543–3557 (2018).

14. Butterfield, G. L. et al. Evolution of a designed protein assembly encapsulating its own RNA genome. Nature 552, 415–420 (2017).

15. Malay, A. D. et al. An ultra-stable gold-coordinated protein cage displaying reversible assembly. Nature 569, 438–442 (2019).

16. Swift, J., Butts, C. A., Cheung-Lau, J., Yerubandi, V. & Dmochowski, I. J. Efficient Self-Assembly of Archaeoglobus fulgidus ferritin around metallic cores. Langmuir 25, 5219–5225 (2009).

17. Sana, B., Johnson, E. & Lim, S. The unique self-assembly/disassembly property of Archaeoglobus fulgidus ferritin and its implications on molecular release from the protein cage. Biochim. Biophys. Acta - Gen. Subj. 1850, 2544–2551 (2015).

18. Nasrollahi, F. et al. Incorporation of Graphene Quantum Dots, Iron, and Doxorubicin in/on Ferritin Nanocages for Bimodal Imaging and Drug Delivery. Adv. Ther. 3, 1900183 (2020).

19. Fletcher, J. M. et al. Self-assembling cages from coiled-coil peptide modules. Science (80-.). 340, 595–599 (2013).

20. Peng, T. & Lim, S. Trimer-based design of pH-responsive protein cage results in soluble disassembled structures. Biomacromolecules 12, 3131–3138 (2011).

21. Dalmau, M., Lim, S. & Wang, S. W. Design of a pH-dependent molecular switch in a caged protein platform. Nano Lett. 9, 160–166 (2009).

22. Pontillo, N., Pane, F., Messori, L., Amoresano, A. & Merlino, A. Cisplatin encapsulation within a ferritin nanocage: A high-resolution crystallographic study. Chem. Commun. 52, 4136–4139 (2016).

23. Liu, X. et al. Apoferritin-CeO2nano-truffle that has excellent artificial redox enzyme activity. Chem. Commun. 48, 3155–3157 (2012).

24. Skouri-Panet, F., Quevillon-Cheruel, S., Michiel, M., Tardieu, A. & Finet, S. sHSPs under temperature and pressure: The opposite behaviour of lens alpha-crystallins and yeast HSP26. Biochim. Biophys. Acta - Proteins Proteomics 1764, 372–383 (2006).

25. White, H. E. et al. Multiple Distinct Assemblies Reveal Conformational Flexibility in the Small Heat Shock Protein Hsp26. Structure 14, 1197–1204 (2006).

26. Zhang, T. et al. Effect of high hydrostatic pressure (HHP) on structure and activity of phytoferritin. Food Chem. 130, 273–278 (2012).

27. Wang, Q. et al. High hydrostatic pressure encapsulation of doxorubicin in ferritin nanocages with enhanced efficiency. J. Biotechnol. 254, 34–42 (2017).

28. Brooks, N. J. et al. Automated high pressure cell for pressure jump x-ray diffraction. Rev. Sci. Instrum. 81, 064103 (2010).

29. Tanner, J. J. Empirical power laws for the radii of gyration of protein oligomers. Acta Crystallogr. Sect. D Struct. Biol. 72, 1119–1129 (2016).

30. Svergun, D., Barberato, C. & Koch, M. H. CRYSOL - A program to evaluate X-ray solution scattering of biological macromolecules from atomic coordinates. J. Appl. Crystallogr. 28, 768–773 (1995).

31. Konarev, P. V., Volkov, V. V., Sokolova, A. V., Koch, M. H. J. & Svergun, D. I. PRIMUS: A Windows PC-based system for small-angle scattering data analysis. J. Appl. Crystallogr. 36, 1277–1282 (2003).

32. Krissinel, E. & Henrick, K. Inference of Macromolecular Assemblies from Crystalline State. J. Mol. Biol. 372, 774–797 (2007).

33. Stefanini, S., Vecchini, P. & Chiancone, E. On the Mechanism of Horse Spleen Apoferritin Assembly: A Sedimentation Velocity and Circular Dichroism Study. Biochemistry 26, 1831–1837 (1987).

34. Gerl, M. & Jaenicke, R. Mechanism of the self-assembly of apoferritin from horse spleen - Cross-linking and spectroscopic analysis. Eur. Biophys. J. 15, 103–109 (1987).

35. Banyard, S. H., Stammers, D. K. & Harrison, P. M. Electron density map of apoferritin at 2.8-Å resolution. Nature 271, 282–284 (1978).

36. Sato, D. et al. Ferritin Assembly Revisited: A Time-Resolved Small-Angle X-ray Scattering Study. Biochemistry 55, 287–293 (2016).

37. Kilic, M. a, Spiro, S. & Moore, G. R. Stability of a 24-meric homopolymer: comparative studies of assembly-defective mutants of Rhodobacter capsulatus bacterioferritin and the native protein. Protein Sci. 12, 1663–1674 (2003).

38. Andrews, S. C. et al. Physical, chemical and immunological properties of the bacterioferritins of Escherichia coli, Pseudomonas aeruginosa and Azotobacter vinelandii. Biochim. Biophys. Acta - Protein Struct. Mol. Enzymol. 1078, 111–116 (1991).

39. Wiedemann, C., Bellstedt, P. & Görlach, M. CAPITO - A web server-based analysis and plotting tool for circular dichroism data. Bioinformatics 29, 1750–1757 (2013).

40. Weber, G. Thermodynamics of the association and the pressure dissociation of oligomeric proteins. J. Phys. Chem. 97, 7108–7115 (1993).

41. Atkins, P. & de, P. J. Atkins’ Physical Chemistry, 7th Edition. Oxford Univ. Press (2002).

42. Ward, A. B., Sali, A. & Wilson, I. A. Integrative Structural Biology. Science (80-.). 339, 913–915 (2013).

43. Brooks-Bartlett, J. C. et al. Development of tools to automate quantitative analysis of radiation damage in SAXS experiments. J. Synchrotron Radiat. 24, 63–72 (2017).

44. Kikhney, A. G. & Svergun, D. I. A practical guide to small angle X-ray scattering (SAXS) of flexible and intrinsically disordered proteins. FEBS Lett. 589, 2570–2577 (2015).

45. Sato, D. et al. Electrostatic Repulsion during Ferritin Assembly and Its Screening by Ions. Biochemistry 55, 482–488 (2016).

46. Watt, R. K., Hilton, R. J. & Graff, D. M. Oxido-reduction is not the only mechanism allowing ions to traverse the ferritin protein shell. Biochimica et Biophysica Acta - General Subjects 1800, 745–759 (2010).

47. Bakker, G. R. & Boyer, R. F. Iron incorporation into apoferritin. The role of apoferritin as a ferroxidase. J. Biol. Chem. 261, 13182–13185 (1986).

48. Rui, H., Rivera, M. & Im, W. Protein dynamics and ion traffic in bacterioferritin. Biochemistry 51, 9900–9910 (2012).

49. Kegel, W. K. & van der Schoot, P. Competing Hydrophobic and Screened-Coulomb Interactions in Hepatitis B Virus Capsid Assembly. Biophys. J. 86, 3905–3913 (2004).

50. Silva, J. Pressure Stability of Proteins. Annu. Rev. Phys. Chem. 44, 89–113 (1993).

51. Silva, J. L., Foguel, D. & Royer, C. A. Pressure provides new insights into protein folding, dynamics and structure. Trends Biochem. Sci. 26, 612–618 (2001).

52. Silva, J. L. & Weber, G. Pressure-induced dissociation of brome mosaic virus. J. Mol. Biol. 199, 149–159 (1988).

53. Leimkühler, M., Goldbeck, A., Lechner, M. D. & Witz, J. Conformational changes preceding decapsidation of bromegrass mosaic virus under hydrostatic pressure: A small-angle neutron scattering study. J. Mol. Biol. 296, 1295–1305 (2000).

54. Leimkühler, M. et al. The formation of empty shells upon pressure induced decapsidation of turnip yellow mosaic virus. Arch. Virol. 146, 653–667 (2001).

55. Teale, F. W. J. Cleavage of the haem-protein link by acid methylethylketone. BBA - Biochim. Biophys. Acta 35, 543 (1959).

56. Nayuk, R. & Klaus, H. Formfactors of Hollow and Massive Rectangular Parallelepipeds at Variable Degree of Anisometry. Zeitschrift für Physikalische Chemie 226, 837 (2012).

57. Willies, S. C., Isupov, M. N., Garman, E. F. & Littlechild, J. A. The binding of haem and zinc in the 1.9 Å X-ray structure of Escherichia coli bacterioferritin. J. Biol. Inorg. Chem. 14, 201–207 (2009).

58. Hingorani, K. et al. Photo-oxidation of tyrosine in a bio-engineered bacterioferritin ‘reaction centre’ - A protein model for artificial photosynthesis. Biochim. Biophys. Acta - Bioenerg. 1837, 1821–1834 (2014).

59. Ardejani, M. S., Li, N. X. & Orner, B. P. Stabilization of a protein nanocage through the plugging of a protein-protein interfacial water pocket. Biochemistry 50, 4029–4037 (2011).

60. Wong, S. G. et al. Structural and mechanistic studies of a stabilized subunit dimer variant of Escherichia coli bacterioferritin identify residues required for core formation. J. Biol. Chem. 284, 18873–18881 (2009).

61. Fan, R., Boyle, A. L., Vee, V. C., See, L. N. & Orner, B. P. A helix swapping study of two protein cages. Biochemistry 48, 5623–5630 (2009).

62. Hargrove, M. S. et al. Stability of Myoglobin: A Model for the Folding of Heme Proteins. Biochemistry 33, 11767–11775 (1994).

63. Andrews, S. C. et al. Overproduction, purification and characterization of the bacterioferritin of Escherichia coli and a C-terminally extended variant. Eur. J. Biochem. 213, 329–338 (1993).

64. Basham, M. et al. Data Analysis WorkbeNch (DAWN). J. Synchrotron Radiat. 22, 853–858 (2015).

65. Rambo, R. P. & Tainer, J. A. Accurate assessment of mass, models and resolution by small-angle scattering. Nature 496, 477–481 (2013).

66. Rambo, R. P. & Tainer, J. A. Characterizing flexible and intrinsically unstructured biological macromolecules by SAS using the Porod-Debye law. Biopolymers 95, 559–571 (2011).

67. Franke, D. & Svergun, D. I. DAMMIF, a program for rapid ab-initio shape determination in small-angle scattering. J. Appl. Crystallogr. 42, 342–346 (2009).

68. Konarev, P. V., Petoukhov, M. V. & Svergun, D. I. Rapid automated superposition of shapes and macromolecular models using spherical harmonics. J. Appl. Crystallogr. 49, 953–960 (2016).

69. Hussain, R., Jávorfi, T. & Siligardi, G. Circular dichroism beamline B23 at the Diamond Light Source. J. Synchrotron Radiat. 19, 132–135 (2012).

70. De Val, N., Declercq, J. P., Lim, C. K. & Crichton, R. R. Structural analysis of haemin demetallation by L-chain apoferritins. J. Inorg. Biochem. 112, 77–84 (2012).

71. Chen, C. R. & Makhatadze, G. I. ProteinVolume: calculating molecular van der Waals and void volumes in proteins. BMC Bioinformatics 16, 101 (2015).

72. Demeler, B. et al. Characterization of size, anisotropy, and density heterogeneity of nanoparticles by sedimentation velocity. Anal. Chem. 86, 7688–7695 (2014).

73. Wald, A. & Wolfowitz, J. On a Test Whether Two Samples are from the Same Population. Ann. Math. Stat. 11, 147–162 (1940).

74. Hansen, S. BayesApp: A web site for indirect transformation of small-angle scattering data. J. Appl. Crystallogr. 45, 566–567 (2012).

