## Supplementary data for "Controlling protein nanocage assembly with hydrostatic pressure"

**Table S1. PISA results for ABfr.**

| Protein | Oligomer | Surface area Å$^2$ | Surface area Å$^2$ | $\Delta G^{int}$ (kcal mol$^{-1}$) | $\Delta G^{diss}$ (kcal mol$^{-1}$) |
|---|---|---|---|---|---|
| ABfr | $M_{24}$ | 132140 | 95240 | -946.6 | 279.5 |
| | $M_2$ | 15140 | 2920 | -76.3 | 11.3 |

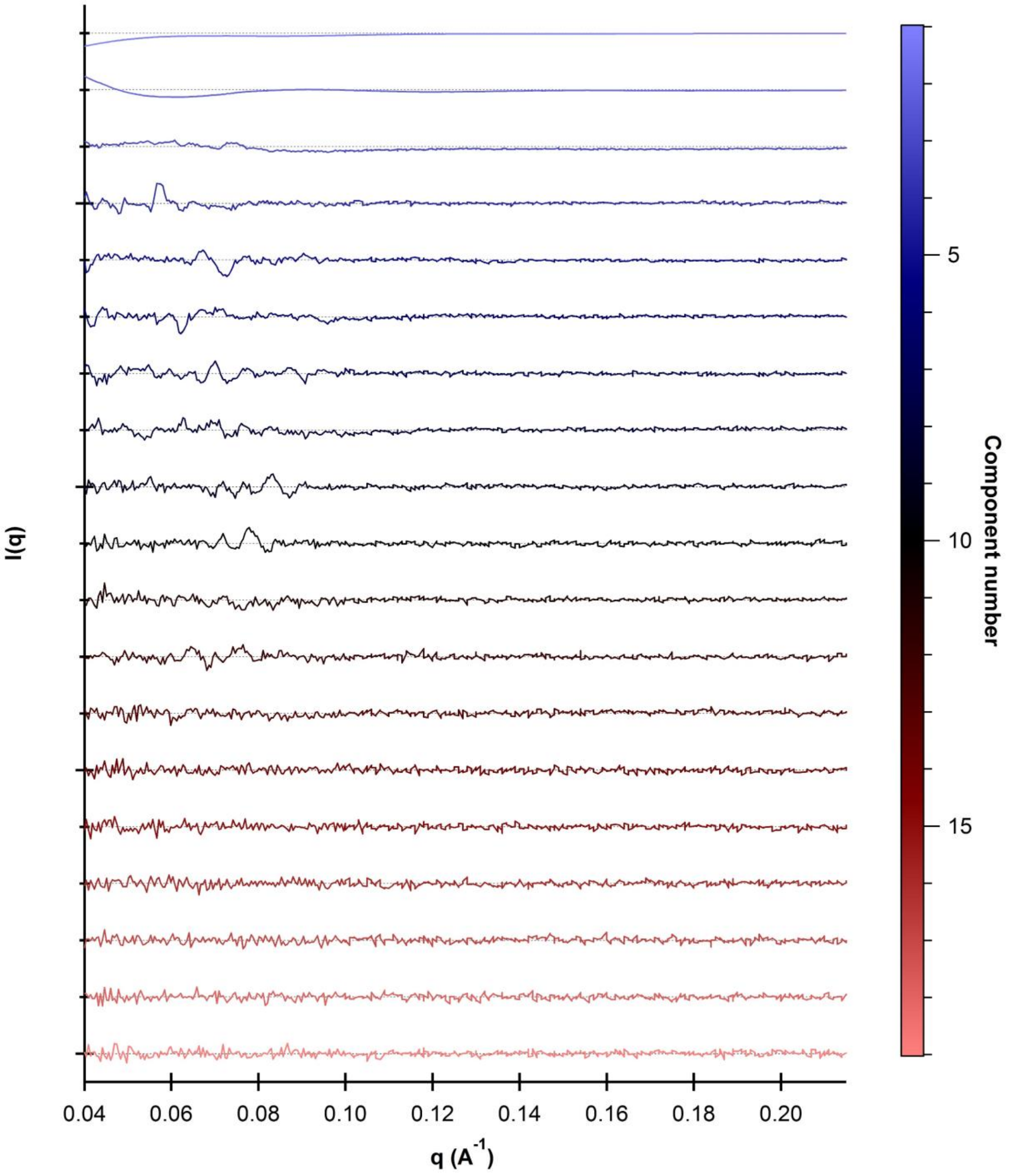


**Figure S1. Eigenvectors for the ABfr dataset, calculated using SVDPLOT.**[1]
Eigenvectors have been offset for clarity, the dotted line represents zero in each case.

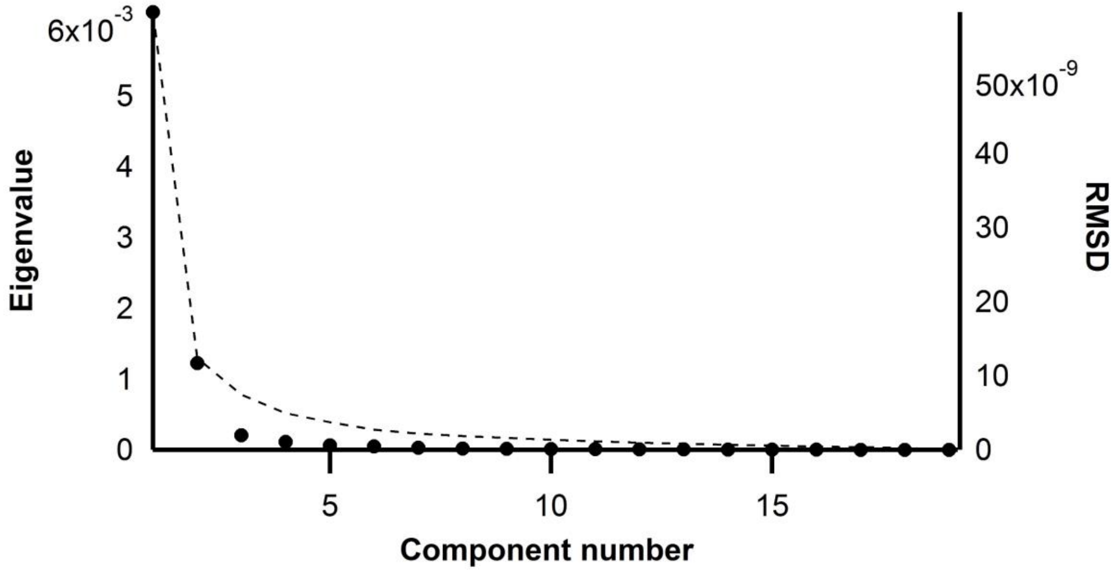


**Figure S2.** Eigenvalues (black circles) for each SVD component and root mean squared deviation for the fit between the reconstructed dataset and experimental dataset with each component incrementally added to the model (dotted line) for ABfr. The first three eigenvalues account for 93.5% of the total.

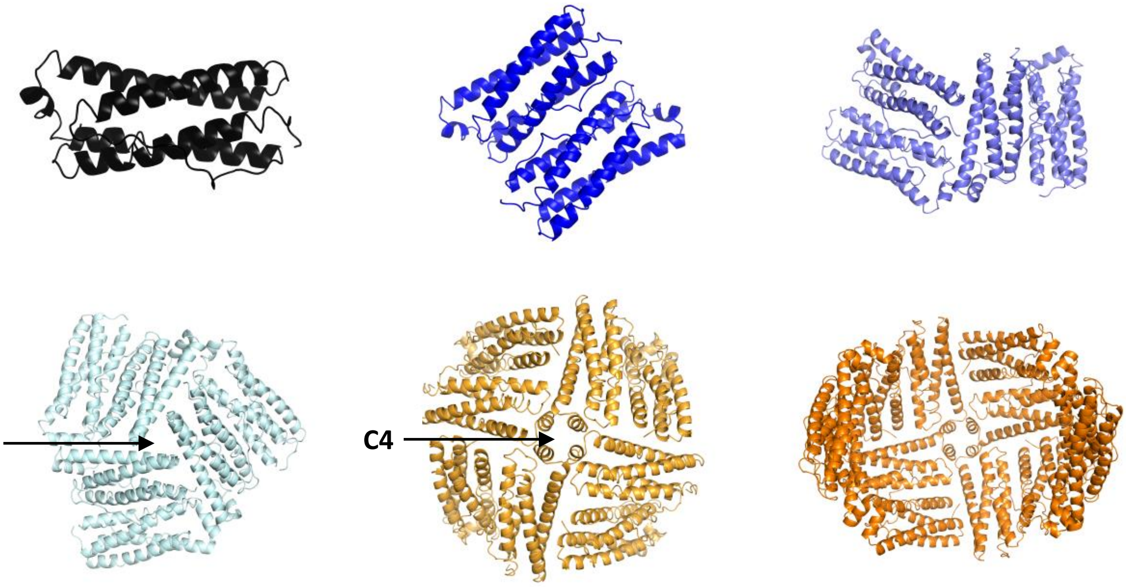


**Figure S3.** Proposed structures for various ABfr subunit oligomers, derived from the full Bfr cage (PBD ID: 2VXI).[2] Relevant symmetry axes are labelled.

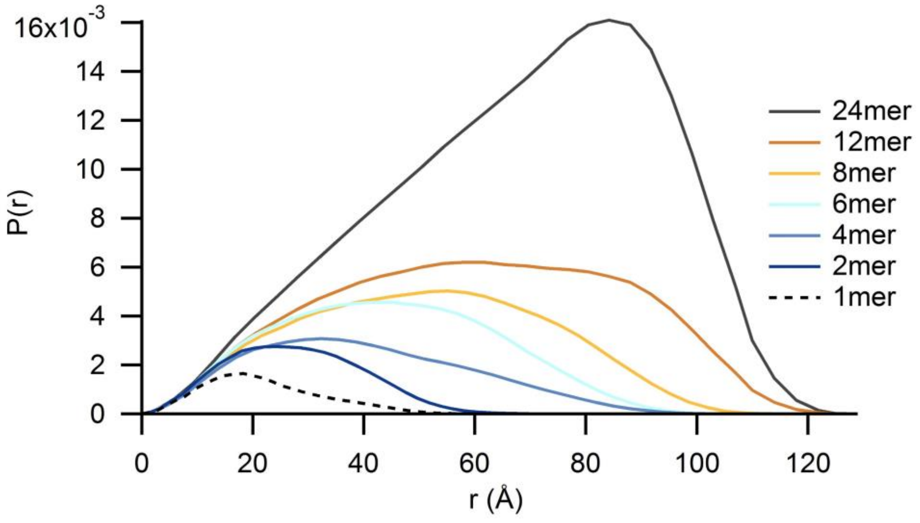


**Figure S4.** P(r) distributions for various ABfr subunit oligomers calculated using CRYSOL and ScÅtter**.**

**Table S2.** Calculated parameters for various Bfr subunit oligomers. P(r) max is the r value at which the P(r) function is maximised.

| Oligomer number | $R_g$ (Å) | Volume (Å$^3$) | $D_{max}$ (Å) | P(r) max (Å) |
|---|---|---|---|---|
| 1 | 17.25 | 20369 | 61 | 18 |
| 2 | 21.37 | 44600 | 74 | 23.4 |
| 4 | 31.24 | 89500 | 108 | 32.4 |
| 6 | 34.4 | 139000 | 108 | 41.7 |
| 8 | 39.51 | 182000 | 117 | 55.2 |
| 12 | 46.77 | 268000 | 131 | 58 |
| 24 | 52.49 | 561000 | 131 | 84.2 |

**Table S3.** Trialled OLIGOMER models and reduced chi-squared values of the global fit.

| Model | Components | Chi-value |
|---|---|---|
| 1 | 24, 1, background | 16.45 |
| 2 | 24, 2, background | 3.78 |
| 3 | 24, 4, background | 7.23 |
| 4 | 24, 6, background | 25.65 |
| 5 | 24, 4, 2, background | 2.66 |
| 6 | 24, 6, 2, background | 2.44 |
| 7 | 24, 6, 4, background | 7.23 |
| 8 | 24, 8, 2, background | 2.24 |
| 9 | 24, 8, 4, background | 7.23 |
| 10 | 24, 12, 2, background | 2.47 |
| 11 | 24, 12, 4, background | 7.23 |

A

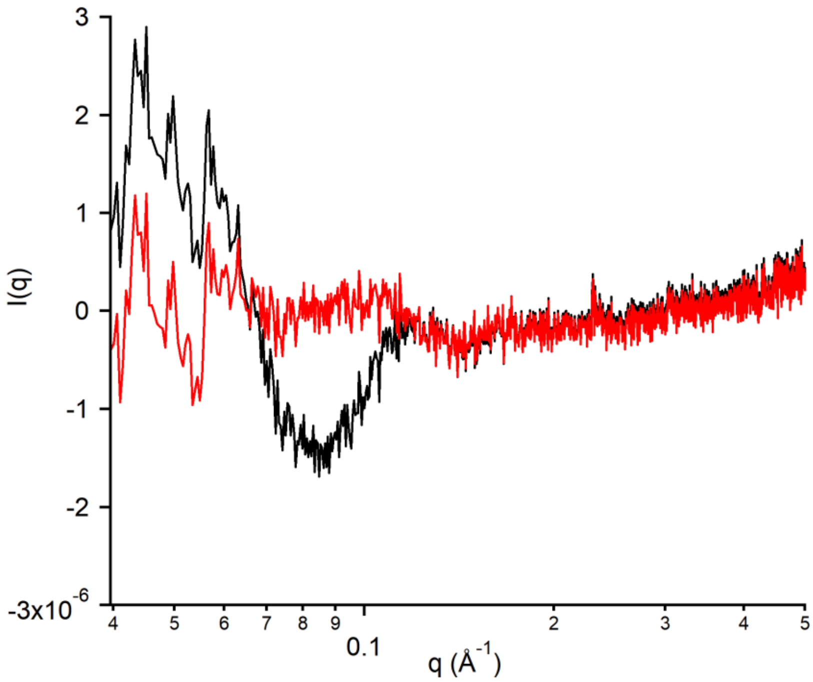


B

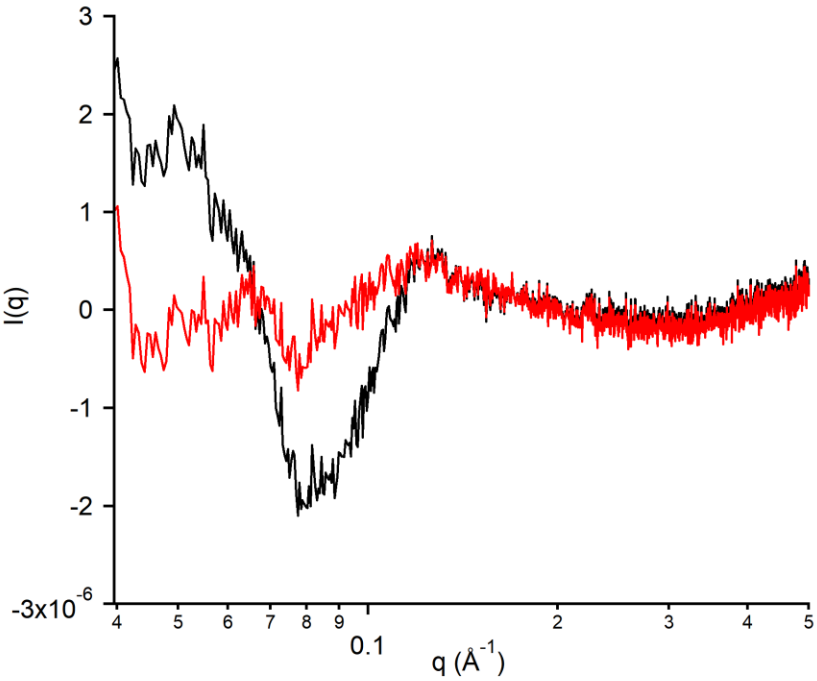


**Figure S5.** Fit residuals of example OLIGOMER models for ABfr at A) 1 MPa and B) 450 MPa. The two-component model (icositetramer + dimer) (black) resulted in poorer fit quality and higher residuals in the low q region than the three component model (icositetramer + hexamer + dimer) (red).

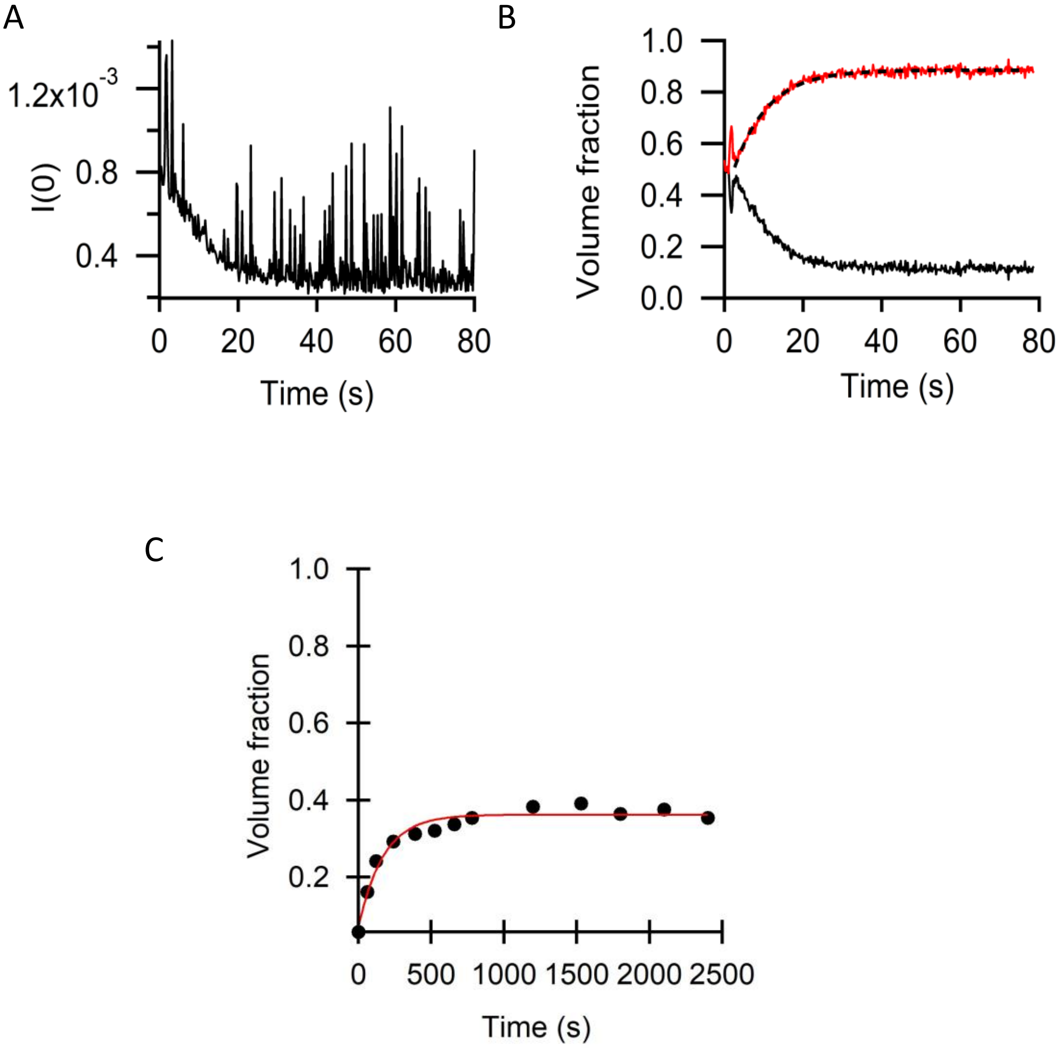


**Figure S6.** ABfr dissociation and reassociation in response to hydrostatic pressure. **(A)** Change in ABfr *I(0)* and **(B)** change in ABfr oligomer volume fraction (dimer – red, icositetramer – black) following rapid pressurisation to 450 MPa. The single exponential fit of dimer fraction is shown as a dotted line, with a rate constant of $k$ = 0.114 ± 0.002 $s^{-1}$. **(C)** Change in ABfr icositetramer volume fraction following pressurization to 450 MPa for 5 minutes. The single exponential fit of icositetramer fraction is shown as a red line, with a rate constant of $k$ = 0.006 ± 0.001 $s^{-1}$.

**Table S4.** AUC data collection parameters and results for ABfr pre- and post- pressurisation.

| **Data collection parameters** | |
|---|---|
| Instrument | Beckman XL-I |
| Wavelength (nm) | 418 |
| Speed (rpm) | 40000 |
| Concentration (μM) | 6.25 |
| Temperature (K) | 293 |
| Buffer density (g $cm^{-3}$) | 1.00276 |
| Buffer viscosity (mPa.s) | 1.021 |

| **Hydrodynamic Parameters** | **Pre-** | **Post-** |
|---|---|---|
| $S_{(20, w)}$ | 15.981 | 16.013 |
| $M_w$ | 446574 | 419223 |
| $f/f_0$ | 1.318 | 1.261 |
| Percentage of total signal | 96% | 97% |
| Stoke’s radius (nm) | 6.67 | 6.24 |

**Table S5. Volume fractions of icositetramer and $R_g$ values for ABfr pre-pressurisation, during pressurisation and post pressure release (t = 1000 s).** Volume fractions are determined from OLIGOMER analysis and so are likely to underestimate the volume fraction of icositetramer.

| | I (mM) | T (˚C) | Initial | | Pressurisation | | End | |
|---|---|---|---|---|---|---|---|---|
| | | | $\Phi_{24}$ | $R_g$ (Å) | $\Phi_{24}$ | $R_g$ (Å) | $\Phi_{24}$ | $R_g$ (Å) |
| **$H_2O$** | 0 | 5 | 0.62 | 49.42 | 0.093 | 30.8 | 0.028 | 29.27 |
| | | 25 | 0.429 | 46.63 | 0.041 | 32.06 | 0.116 | 45.16 |
| **NaPi** | 123 | 5 | 0.465 | 49.12 | 0.105 | 28.04 | 0.099 | 44.07 |
| | | 25 | 0.255 | 49.42 | 0.112 | 39.98 | 0.234 | 49.21 |
| **NaPi + NaCl** | 373 | 5 | 0.734 | 50.36 | 0.059 | 24.45 | 0.298 | 49.06 |
| | | 25 | 0.77 | 51.12 | 0.124 | 35.24 | 0.529 | 49.59 |

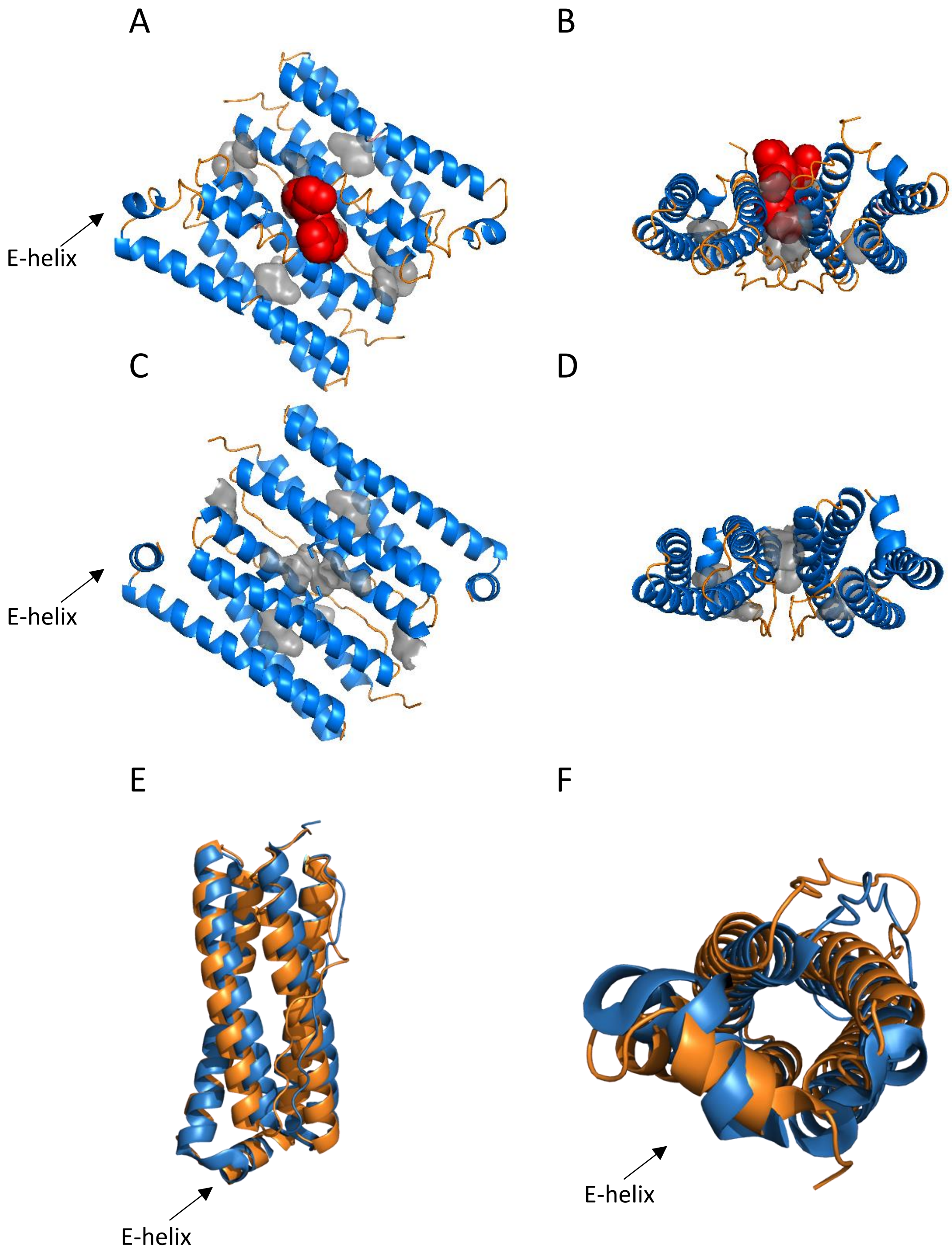


**Figure S7. Crystal structures of the subunit dimer of bacterioferritin.** (A) and (B) show the heme containing protein (ABfr), with distortion of helical structure around the heme binding pocket. (C) and (D) show the apo-protein (AABfr). Alpha helical structure is shown in blue, disordered structure is shown in orange, voids in the structure are shown in grey, and the heme cofactor is shown in red. (E) and (F) show overlaid ABfr (2VXI) (orange) and AABfr (4CVP) (blue) monomer structures[2,3].

A

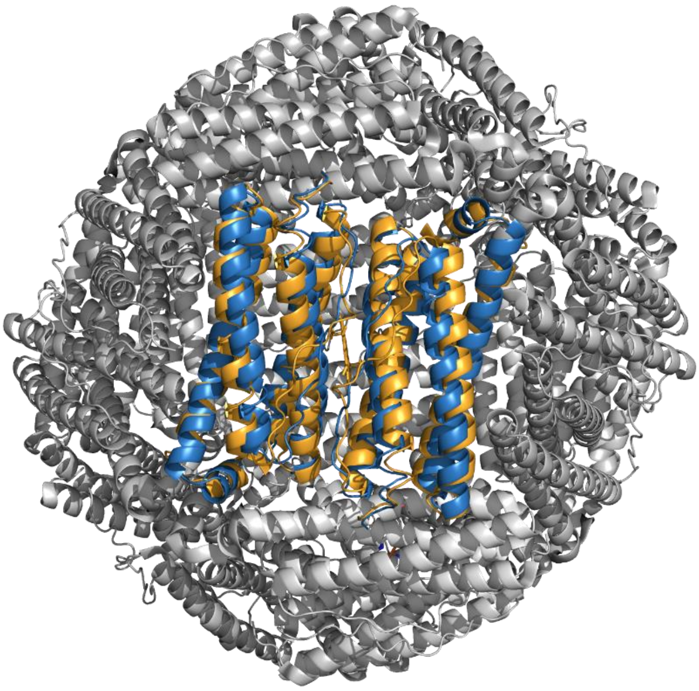

B

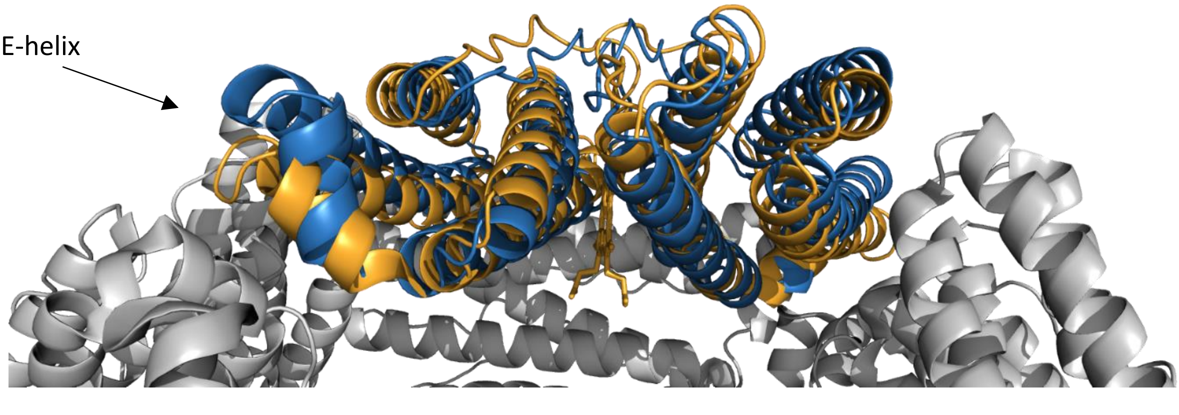


**Figure S8.** (A) ABfr icositetramer (2VXI) (grey / orange) with overlaid AABfr subunit dimer (4CVP) (blue), and (B) cutaway[2,3].

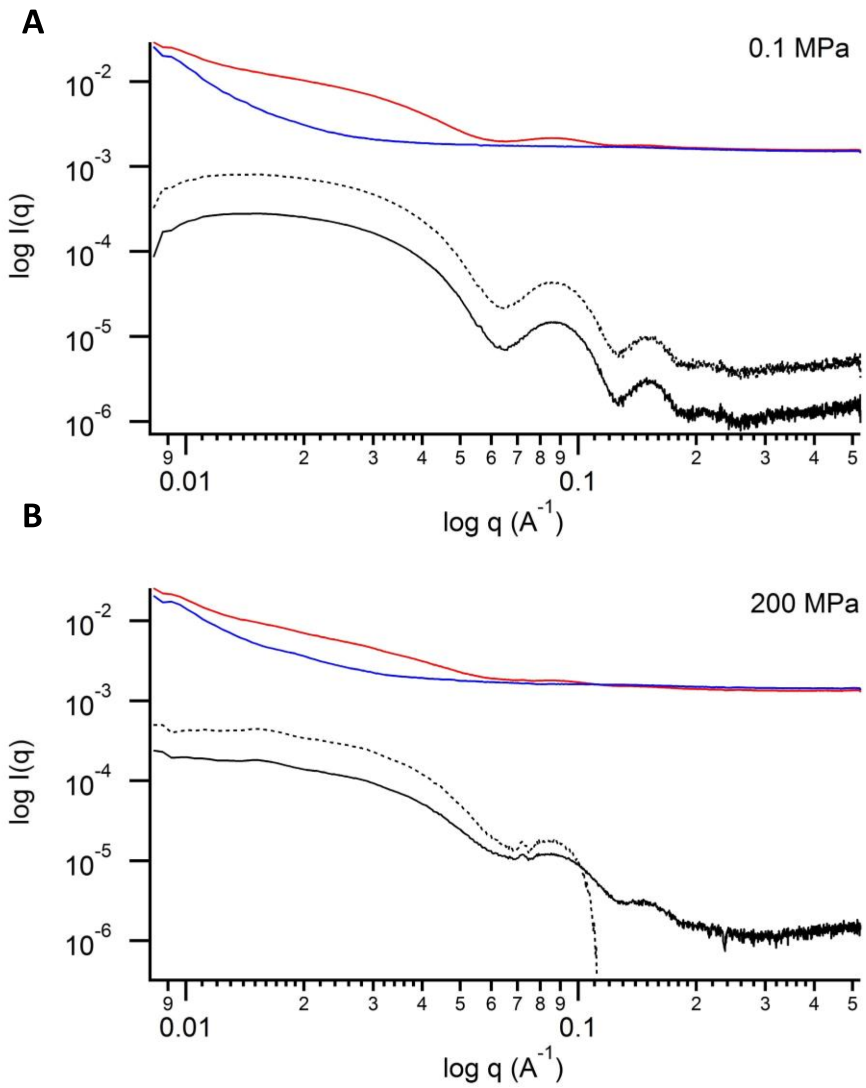


**Figure S9.** Unsubtracted sample data (red) and background data (blue) for ABfr at 0.1 MPa **(A)** and 200 MPa **(B)**. Subtracted sample data sets are shown with background adjustment (dashed line) and without background adjustment (solid line).

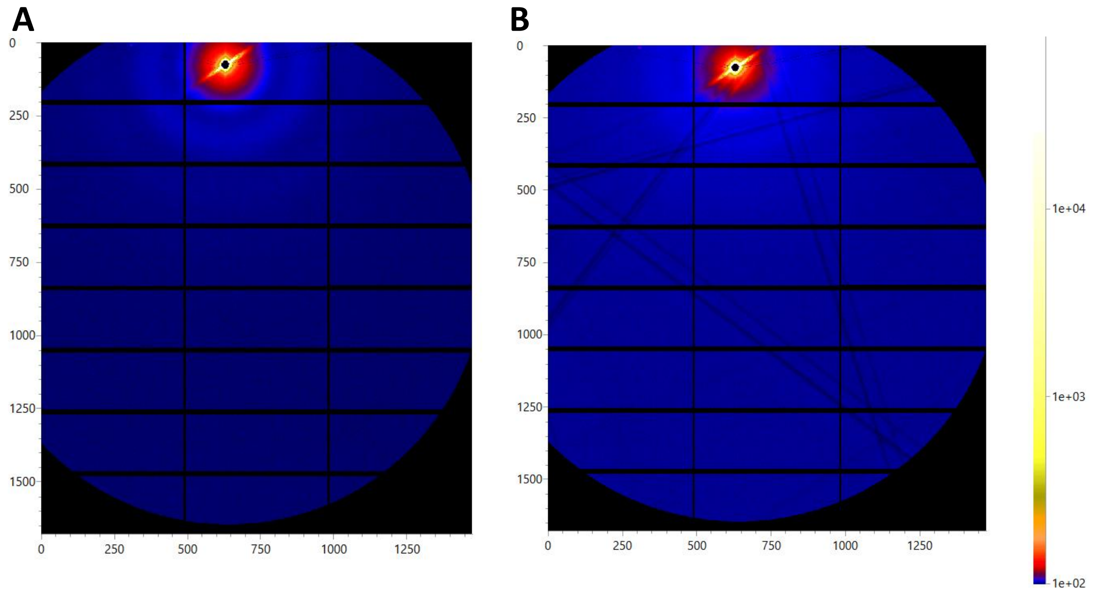


**Figure S10.** 2D detector images (Pilatus P3-2M) for ABfr at 0.1 MPa (A) and 200 MPa (B). Kossel lines are visible at 200 MPa as dark lines and increased intensity and flaring close to the beam-stop.

### References

1. Konarev, P. V., Volkov, V. V. & Svergun, D. I. Interactive graphical system for small-angle scattering analysis of polydisperse systems. *J. Phys. Conf. Ser.* **747**, 012036 (2016).

2. Willies, S. C., Isupov, M. N., Garman, E. F. & Littlechild, J. A. The binding of haem and zinc in the 1.9 Å X-ray structure of Escherichia coli bacterioferritin. *J. Biol. Inorg. Chem.* **14**, 201–207 (2009).

3. Hingorani, K. *et al.* Photo-oxidation of tyrosine in a bio-engineered bacterioferritin 'reaction centre' - A protein model for artificial photosynthesis. *Biochim. Biophys. Acta - Bioenerg.* **1837**, 1821–1834 (2014).
